# The Generation and Propagation of the Human Alpha Rhythm

**DOI:** 10.1101/202564

**Authors:** Mila Halgren, István Ulbert, Hélène Bastuji, Dániel Fabó, Lorand Erőss, Marc Rey, Orrin Devinsky, Werner K. Doyle, Rachel Mak-McCully, Eric Halgren, Lucia Wittner, Patrick Chauvel, Gary Heit, Emad Eskandar, Arnold Mandell, Sydney S. Cash

## Abstract

The alpha rhythm is the longest studied brain oscillation and has been theorized to play a key role in cognition. Still, its physiology is poorly understood. In this study, we used micro and macro electrodes in surgical epilepsy patients to measure the intracortical and thalamic generators of the alpha rhythm during quiet wakefulness. We first found that alpha in posterior cortex propagates from higher-order anterosuperior areas towards the occipital pole, consistent with alpha effecting top-down processing. This cortical alpha leads pulvinar alpha, complicating prevailing theories of a thalamic pacemaker. Finally, alpha is dominated by currents and firing in supragranular cortical layers. Together, these results suggest that the alpha rhythm likely reflects short-range supragranular feedback which propagates from higher to lower-order cortex and cortex to thalamus. These physiological insights suggest how alpha could mediate feedback throughout the thalamocortical system.

## Main Text

Alpha oscillations (7-13 Hz)^1^ are the most salient EEG event during wakefulness and may be fundamental for top-down cognitive processes^2,3^ such as attention^4^, perception^5,6^, functional inhibition^7^ and working memory^8^. However, the underlying neural structure(s) and circuits which generate alpha are intensely controversial. Studies have pointed to the thalamus as the primary alpha pacemaker, with the classic posterior alpha rhythm driven by the pulvinar and/or lateral geniculate nucleus (LGN)^4,9–11^. Within the cortex, it’s widely assumed that alpha originates from infragranular layers driven by layer V pyramidal cells^12–16^. Despite the prevalence of these hypotheses, the studies used to support them are not definitive; previous electrophysiological literature have either used a distant reference susceptible to volume conduction^4,12,14^, were performed in vitro^13^ or relied on extracranial recordings^17^ (see **Discussion**). Crucially, none of these hypotheses have been directly tested via invasive recordings in humans. We therefore analyzed focal micro and macro electrode recordings from human neocortex and thalamus in surgical epilepsy patients to characterize alpha’s generation during quiet wakefulness.

We analyzed electrocorticography (ECoG) recordings of spontaneous alpha oscillations (4.54±.87 minutes, mean ± standard deviation) in the occipital, posterior temporal, and parietal cortices of 5 patients (3 of whom performed an eye closure task) (**Supplementary Fig. 1, Supplementary Table 1**) (ECoG Patients E1-5). Strikingly, alpha oscillations propagated as travelling waves from anterosuperior cortex towards posteroinferior areas (**Fig. 1-2, Supplementary Fig. 4**)^18^. To quantify this propagation, we used a two-pass third-order zero-phase shift Butterworth Filter between 7-13 Hz to extract alpha-band activity. The Hilbert Transform was then applied to find both the amplitude and phase of ongoing alpha activity, and only timepoints with the highest 20% of alpha-band amplitude (averaged across array channels at each timepoint) were analyzed further.

**Fig 1:**
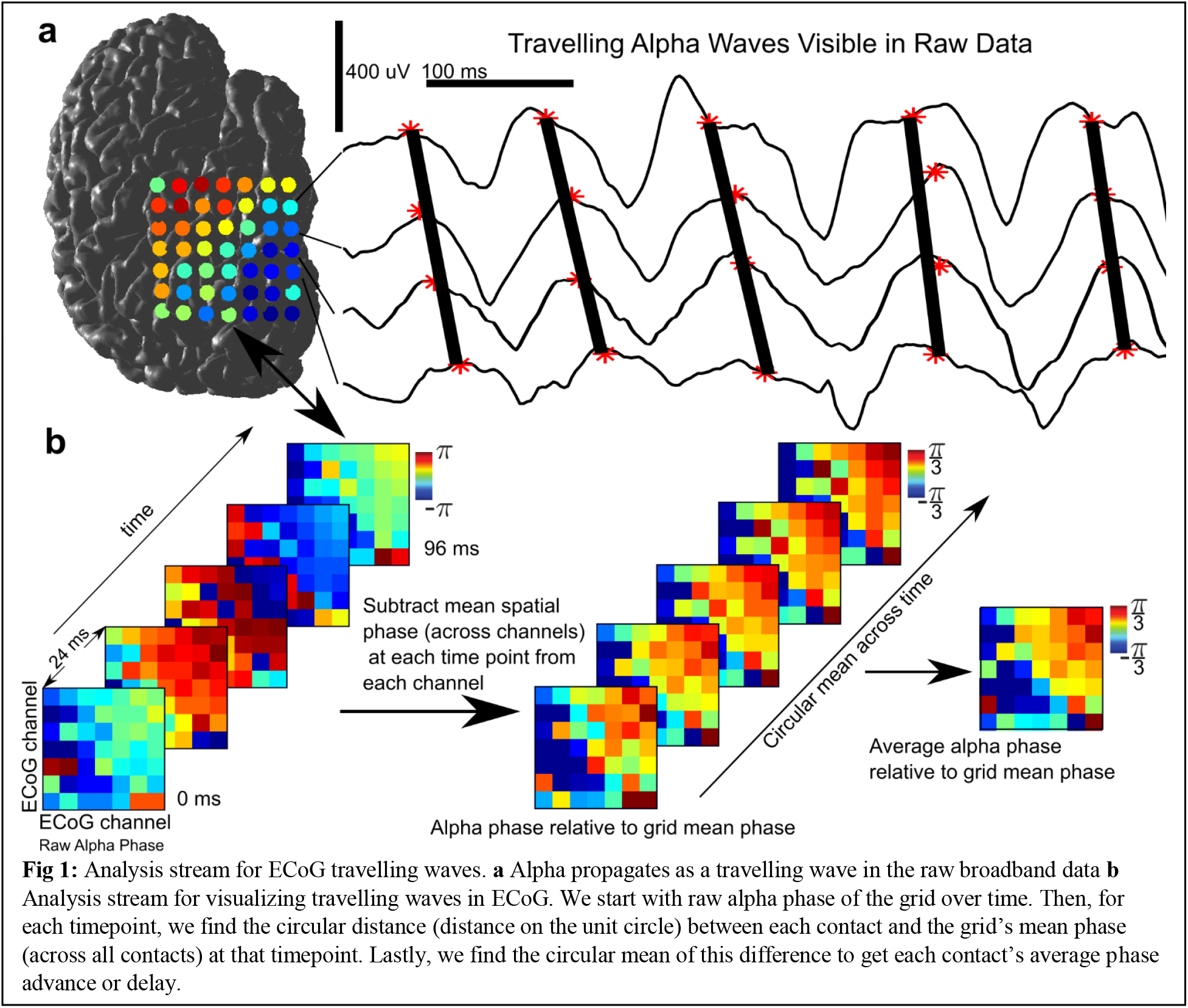
Analysis stream for ECoG travelling waves. **a** Alpha propagates as a travelling wave in the raw broadband data **b** Analysis stream for visualizing travelling waves in ECoG. We start with raw alpha phase of the grid over time. Then, for each timepoint, we find the circular distance (distance on the unit circle) between each contact and the grid’s mean phase (across all contacts) at that timepoint. Lastly, we find the circular mean of this difference to get each contact’s average phase advance or delay.

**Fig. 2:**
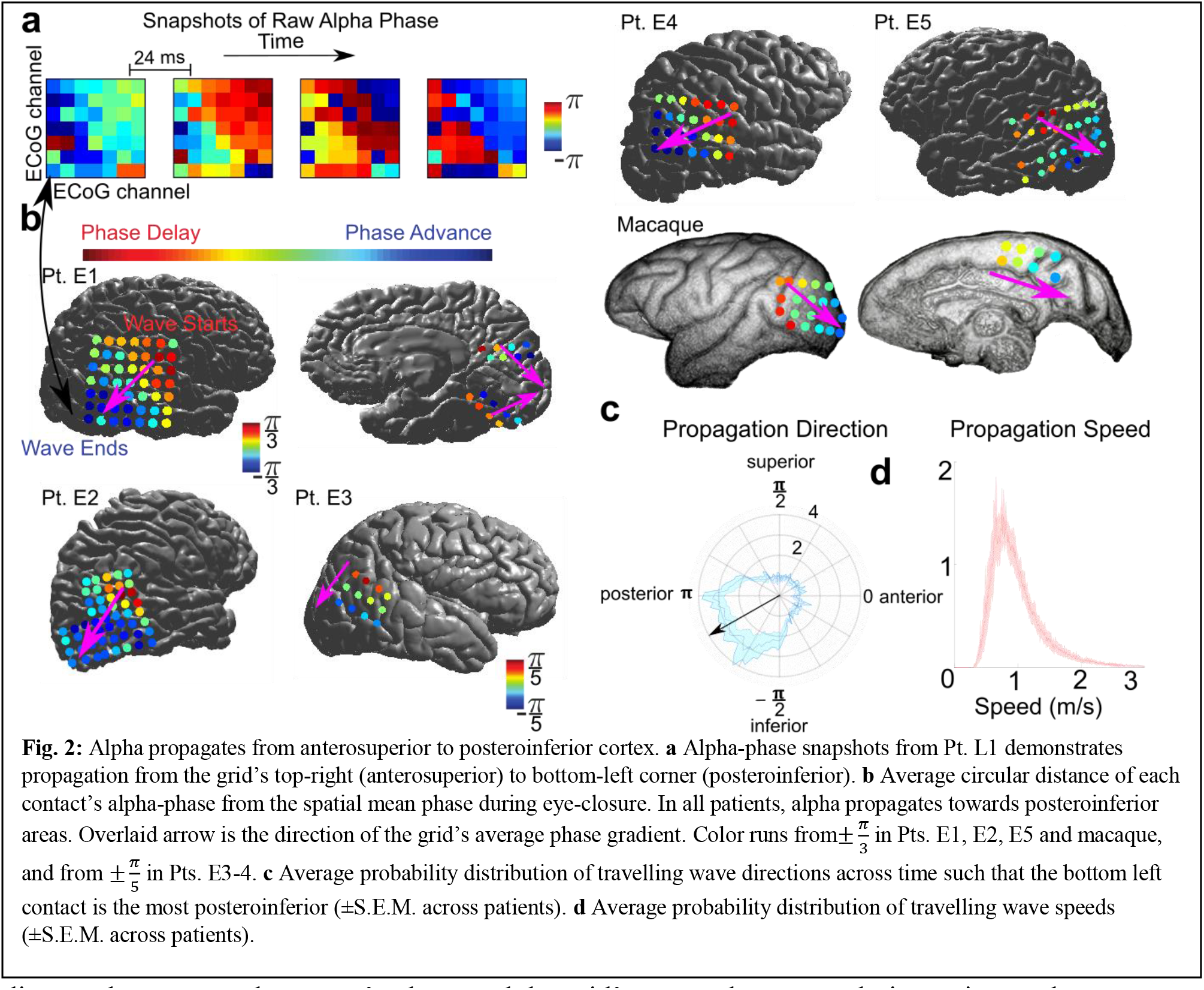
Alpha propagates from anterosuperior to posteroinferior cortex. **a** Alpha-phase snapshots from Pt. L1 demonstrates propagation from the grid–s top-right (anterosuperior) to bottom-left corner (posteroinferior). **b** Average circular distance of each contact’s alpha-phase from the spatial mean phase during eye-closure. In all patients, alpha propagates towards posteroinferior areas. Overlaid arrow is the direction of the grid’s average phase gradient. Color runs from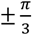 in Pts. E1, E2, E5 and macaque, and from 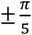 in Pts. E3-4. **c** Average probability distribution of travelling wave directions across time such that the bottom left contact is the most posteroinferior (±S.E.M. across patients). **d** Average probability distribution of travelling wave speeds (±S.E.M. across patien

To visualize the spatial progression of alpha oscillations across the array, we found the circular difference between the mean phase across all contacts and each individual contact at each point in time (**Fig. 1b**). This yields a distribution of differences of each contact’s phase from the grid’s mean phase across all time-points. We then found the circular mean of this difference: if a contact is leading an oscillation, it will have a positive circular distance with respect to the grid’s spatial mean phase; if a contact is lagging, it will have a negative phase difference with the grid’s average phase. **Fig. 2b** was then generated by finding the mean circular distance between each contact’s phase and the grid’s mean phase at each timepoint, or the average advance/delay of a given contact. This method allows one to measure travelling waves oblique to the grid’s implantation and sidestep the selection of a potentially biasing reference contact. We confirmed that alpha had a consistent propagation direction in each patient by finding the direction of the average spatial phase gradient across the grid of electrodes at each time point, and then determining if the distribution of gradient directions throughout the grid was non-uniform^19^ (p ≤ 10^−17^ in each patient, Rayleigh Test) (**Fig. 2d, Supplementary Fig. 3, Methods**). Estimated median speeds of these waves (derived from the phase-gradient) were just under 1 m/s (median speed across patients: .9134 +-.1563 m/s). (**Supplementary Fig. 4, Methods**). Open-source ECoG recordings in a healthy macaque during eye closure demonstrated a highly similar propagation direction and speed^20,21^ (**Fig. 2b**).

To determine if the thalamus coordinated these travelling alpha waves, we utilized S(tereo)EEG to make bipolar local-field-potential gradient (LFPg) macroelectrode depth recordings (n=9 patients, 36 ± 7.5 minutes, mean±standard deviation) during quiet wakefulness.Recordings were made simultaneously from cortex and the pulvinar, a thalamic nucleus which projects broadly to posterior cortical regions^22^ and postulated to drive cortical alpha^4,23^ (**Fig. 3a**) (SEEG Patients S1-9). The use of a bipolar derivation (i.e. referencing each contact to its neighbor) ensured that activity was locally generated, and not volume conducted from a distal structure. Cortical coverage was predominantly posterior (108/124 cortical contacts posterior to the central sulcus), similar to our ECoG patients (**Fig. 3b**). We first verified that alpha travelling waves could be measured in these cortical depth recordings by applying the same method used to quantify alpha propagation in our ECoG data (i.e. measuring how much each individual channel’s alpha phase lead or lagged the mean phase across all channels). Analysis was restricted to occipital and posterior temporal/parietal channels which were in unambiguously lateral cortex to avoid spurious phase inversions, and (just as with ECoG) to time-points with the top 20% of cortical alpha power (averaged across channels at each timepoint). The alpha phase of anterosuperior contacts led ones closer to the occipital pole, replicating our ECoG recordings (**Fig. 4a**). This demonstrates that these travelling waves are not reference-dependent and suggests that the alpha rhythms recorded in our depth patients are analogous to the ones recorded in our ECoG patients. As we then wished to examine the pulvinar’s role in cortical alpha, all further analyses were biased towards thalamic activity by only analyzing the two-second epochs with the 20% most thalamic alpha-band power (averaged across all thalamic channels). We first wished to characterize the prevalence of alpha rhythms in both cortex and pulvinar. This was done by detecting which channels had peaks between 7-13 Hz in their power spectra (peaks were detected via the peakfinder algorithm; see **Methods**). Surprisingly, power spectra from cortical contacts had alpha-band peaks more frequently (63.4%, 78/123 of cortical channels) than ones in the pulvinar (34.6%, 9/26 of thalamic channels) (**Fig. 4b, Supplementary Fig. 6, Methods**) Thalamic and cortical power spectra also sometimes had different peak frequencies (**Fig. 4b**); while this could be construed as evidence for separate thalamic and cortical alpha generators, this is not necessarily the case. Empirically, spindles (believed to be thalamocortically driven) have higher frequencies in the thalamus than the cortex^24^. Analytically, weakly coupled oscillators can also exhibit different peak frequencies despite driving one another^25^. Thalamocortical coherence spectra often exhibited robust alpha-peaks (**Fig. 4e**), indicating that alpha rhythms in posterior cortex and pulvinar are functionally coupled (peak alpha coherence in thalamocortical channel pairs with significant alpha coherence: .3346+-.012, mean+-s.e.m.). As a first means of determining whether neocortical alpha led thalamic alpha (or vice-versa), we measured the cross-covariance of alpha-amplitude (as derived from the amplitude envelope Hilbert transform) in thalamocortical channel pairs with statistically significant alpha coherence^26^. Though a slight majority of cross-covariances had peaks indicating cortical alpha led thalamic alpha, this was not significant (cortex led thalamus in 70/128 of the thalamocortical channel pairs with a non-zero cross-covariance peak), and the average cross-covariance peaked at zero (**Fig. 4c**). However, the integrated area under the cross-covariance curve between 0 and 400 ms for all thalamocortical channel pairs indicated a cortical lead in the onset of alpha power. This could be seen in both the average cross-covariance (**Fig, 4c**), and quantified on the single-channel level by comparing the power under the curve for cortex leading thalamus vs. thalamus leading cortex (Wilcoxon sign rank, p=2.557 * 10^−4^).

**Fig. 3.**
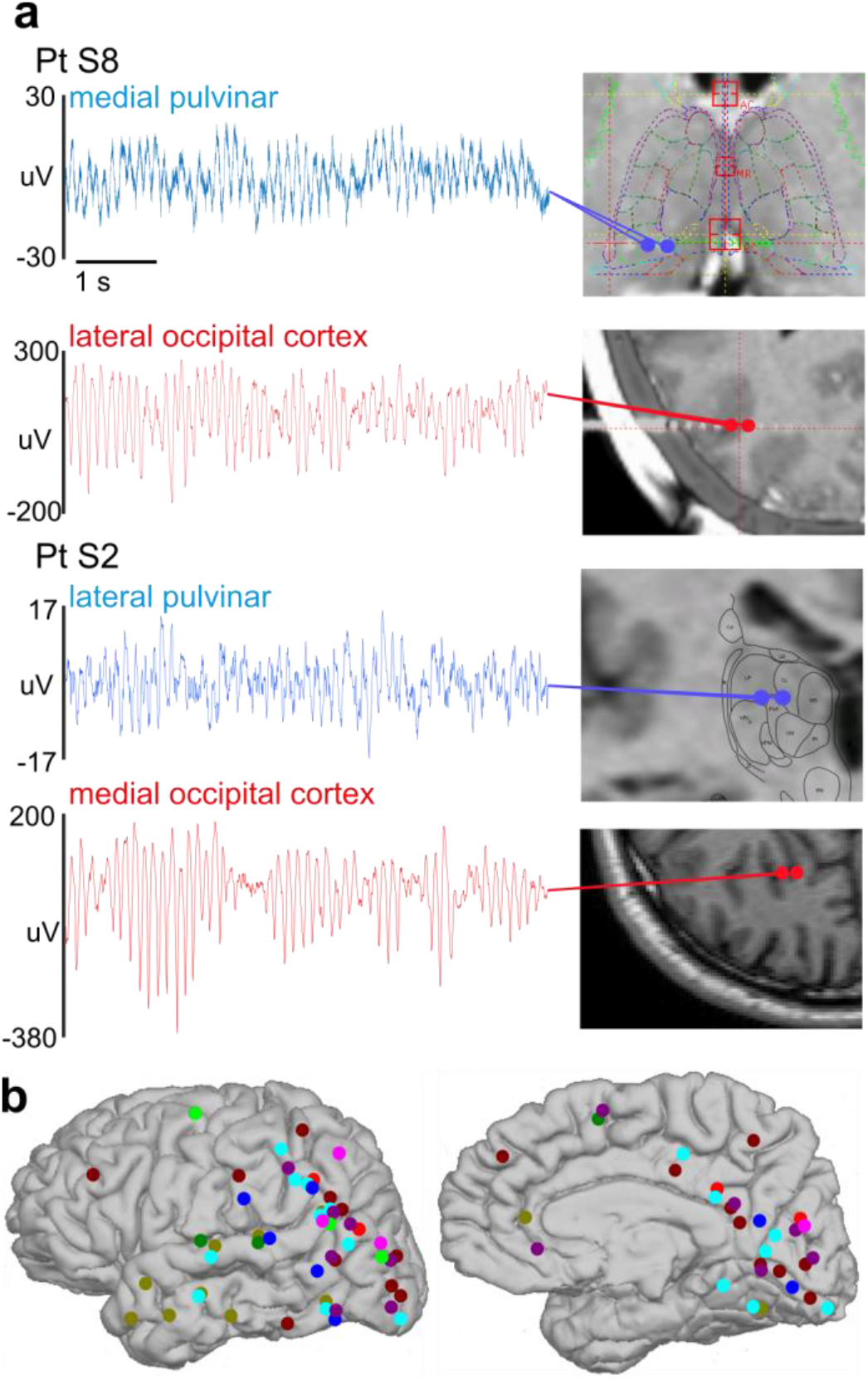
: Robust alpha rhythms can be recorded in human pulvinar and cortex. **a** Representative 6-second traces of simultaneous thalamic and cortical alpha activity. Prominent, largely continuous alpha rhythms can be recorded in various locations within the pulvinar as well as posterior cortex. **b** Cortical implant locations in all SEEG patients displayed on Pt. S3’s brain. Each color signifies a different patient.

**Fig. 4.**
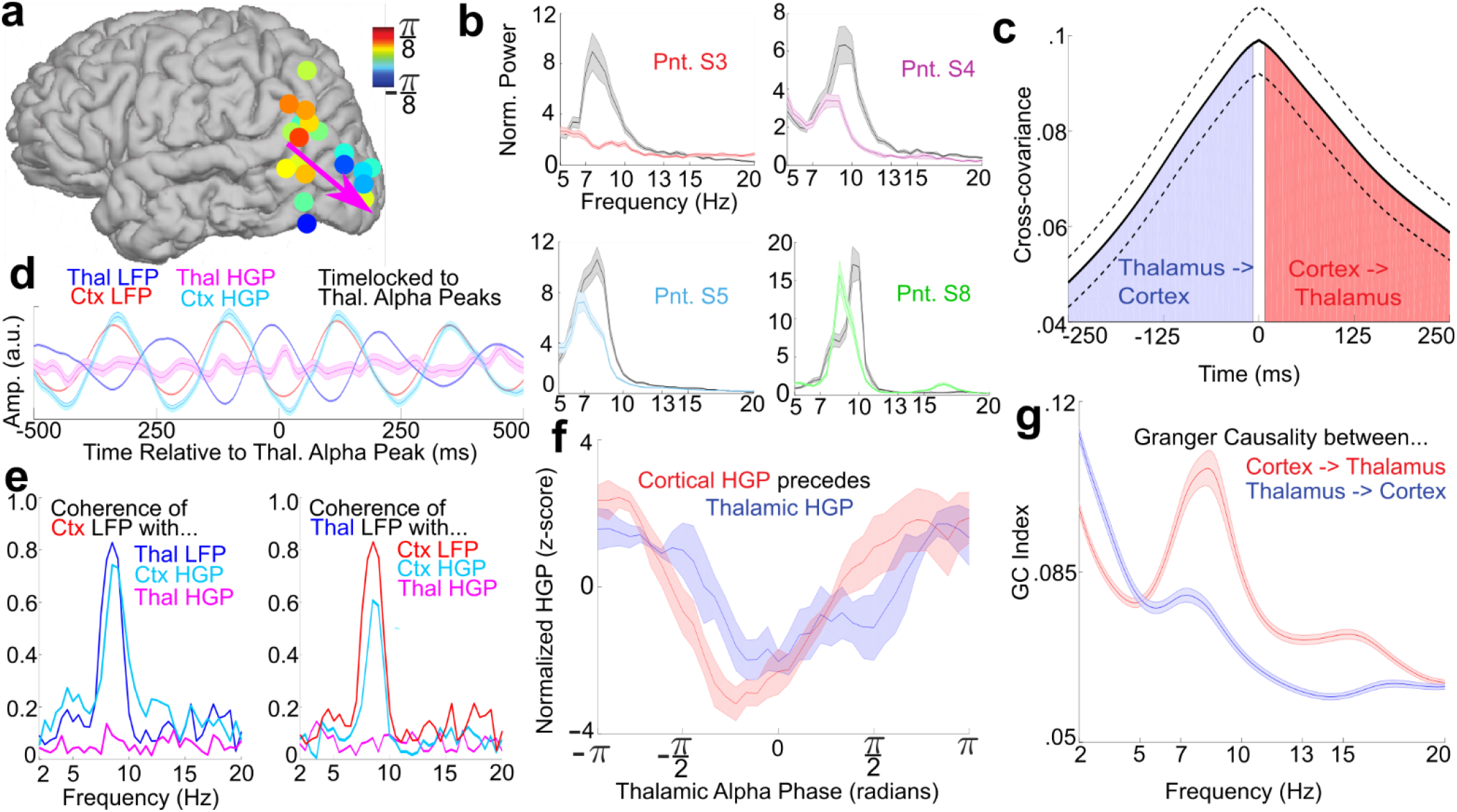
: Cortical alpha leads thalamic (pulvinar) alpha. **a** Average alpha phase lag/leads in bipolar contacts. Note that anterosuperior channels lead inferposterior ones, in accord with our ECoG recordings. **b** Power spectra of the thalamic (color) and cortical (grey) channel with the greatest alpha power. **c** Average cross-covariance between thalamic and cortical alpha power; though the cross-covariance peaks at 0 ms on average, there is significantly more cross-covariance power for cortex leading thalamus than thalamus leading cortex. **d** Cortical and thalamic LFPg and high-gamma-power (HGP) from representative channels locked to peaks in thalamic alpha LFPg-cortical, but not thalamic, firing is phasic with thalamic LFPg. **e** Coherence spectra of thalamic and cortical LFPg with thalamic and cortical HGP and LFPg from the same channel pair in **d**; the coherence of thalamic LFPg with cortical (but not thalamic) suggests that the cortex may drive thalamic alpha activity. **f** Normalized thalamic and cortical HGP at different thalamic alpha phases averaged across channels; note that cortical HGP slightly leads thalamic HGP. **g** Granger Causality spectra averaged over all thalamocortical contact pairs corticothalamic causality shows a strong alpha peak.

To determine whether cortical or thalamic activity was driving these rhythms, we extracted high-gamma-power (HGP), a proxy for neural firing, in both structures (n=5; 70-120 Hz in Pts. S1-3 due to a low sampling rate, 70-190 Hz in Pts. S8-9, Pts. S4-7 were excluded due to low sampling rates; note that using the same 70-120 Hz HGP band in S8-9 didn’t substantially change the results, **Methods**). If a given structure is generating alpha oscillations (and if local HGP reflects neural firing), its HGP should be synchronous with its alpha-band LFPg and exhibit phase-amplitude-coupling (PAC). PAC was assessed using two methods; Tort’s Modulation Index (MI)^27^ and the coherence between the time-domain LFPg and HGP ^28^ (**Fig. 4e, Supplementary Fig. 7**). To ensure that this PAC wasn’t spuriously driven by sharp waveforms^29^, we measured the Sharpness Ratio (SR, see **Methods**) of each channel’s alpha, and then measured the correlation of this with the strength of each channel’s PAC quantified by its MI. SR and MI were not significantly correlated, trending towards anticorrelation (i.e. smoother waveforms had more PAC), the opposite of what would be expected if our PAC was spurious (r=-.2124, p=.0602).^29,30^ Notably, thalamic alpha was rarely coherent with its own HGP (Coherence: 0/14 intrathalamic contact pairs, MI: 3/14, p < .05 Bonferroni Corrected within patients); Instead, thalamic alpha rhythms were predominantly synchronous with cortical HGP (Coherence: 9/14, MI:10/14; mean peak alpha coherence between thalamic LFPg and cortical HGP channel pairs with significant coherence of .3535+-.026) (**Fig. 4e**), supporting cortical generation.

Because HGP is an imperfect proxy for neuronal firing^31^, the failure to find consistent local thalamic PAC could reflect a limitation of our recordings rather than a cortical origin for alpha (but it should be noted that thalamic HGP is modulated by thalamic sleep spindles in the same recordings^24^). To resolve this ambiguity, we further analyzed the minority of thalamic channels (5/42 intrathalamic contact pairs) in which alpha LFPg was phasic with HGP in at least one thalamic and cortical channel. These channel pairs gave us the opportunity to examine average HGP in both thalamus and cortex at different thalamic alpha phases. Unlike relative LFPg phase, which is uninterpretable in the thalamus due to its non-laminar structure, differences in mean HGP with respect to alpha LFPg phase can be interpreted as lags of population activity^24^. Averaging the mean cortical and thalamic HGP with respect to the phase of thalamic alpha LFPg phases across channel pairs, it is apparent that cortical HGP leads thalamic HGP (**Fig 4f**). In individual thalamocortical channel pairs, this could be quantified by measuring the cross-covariance between the thalamic and cortical HGP profiles and seeing if the peak was positive/negative, or examining which had minimal HGP at an earlier thalamic alpha phase.Cortex led thalamus in all 5 channel pairs as derived by both measures, more than expected by chance (p=.0313, one-tailed binomial test of cortex leading thalamus against thalamus leading cortex) (**Fig. 4f**). This lag (difference between HGP minima, as seen in **Fig. 4f**) was on average ~40 degrees, or ~11ms assuming an alpha period of 100 ms. This time delay is physiologically plausible and similar to how much the thalamus leads cortex during sleep spindles^24^.

As a final directional measure, we measured the Granger Causality spectrum (which quantifies the amount of information one time-series contains about another across frequencies) of the LFPg between all pairs of cortical and thalamic contacts^32^ (**Fig. 4g**). Corticothalamic causality in the alpha band was found to be significantly greater than thalamocortical causation for almost every thalamocortical channel pair (across all patients) with a significant difference between thalamocortical and corticothalamic causation (p ≤.01 for each channel pair, Wilcoxon Signed Rank Test, Bonferroni Corrected within patients; 143/163 (87.73%) pairs with greater corticothalamic than thalamocortical causality; p < 1.83 * 10^−24^ across all significantly different channel pairs, Binomial Test). To ensure that this wasn’tdue to our cortical channels having greater alpha power, we repeated our Granger analysis only using thalamocortical channel pairs in which the thalamic lead had greater normalized alpha power. This actually increased the percentage of channel pairs with significantly greater corticothalamic than thalamocortical alpha causality (74/82 (90.24%), significantly more than chance as determined by the binomial test, p < 7.2 * 10^−15^).

To determine which cortical layers generate the alpha rhythm, we utilized laminar microelectrodes^33^ in occipital, temporal and parietal cortex to record current-source-density (CSD, n=3), HGP and Multiunit activity (MUA) (n=2) across gray matter layers during quiet wakefulness (11.32 ± 0.48 minutes, mean ± standard deviation)^33^ (**Fig. 5**) (Laminar Patients L1-3). The CSD is the second spatial derivative of the monopolar field potential, which yields a volume-conduction free measure of local transmembrane currents surrounding the laminar probe^33,34^. MUA (filtered online at 200-5000 Hz, then filtered offline at 300-3000 Hz and rectified^33^) and HGP (filtered offline at 70-190 Hz) are also spatially focal and reflect neural firing^35^. By quantifying both transmembrane currents (which generally reflect postsynaptic events^36^) and firing within each cortical layer, we can determine which laminae generate the alpha-currents and firing measured extracortically with ECoG, MEG and EEG^34,37–39^. Similar to our previous analyses, we only utilized artifact-free two-second epochs with the 20% most alpha-band power. Despite being recorded from various regions of cortex, alpha-band currents in all patients were strongest within superficial cortical layers (Pt. L1: p<2.27 * 10^−25^, Pt. L2: p<1.9 * 10^−22^, Pt. L3: p<1.66 * 10^−5^; largest p-values of Wilcoxon Sign Rank comparing mean alpha power in supragranular versus granular and infragranular channels across epochs, Bonferroni corrected) (**Fig. 6b, e, Supplementary Fig. 8**).

**Fig. 5:**
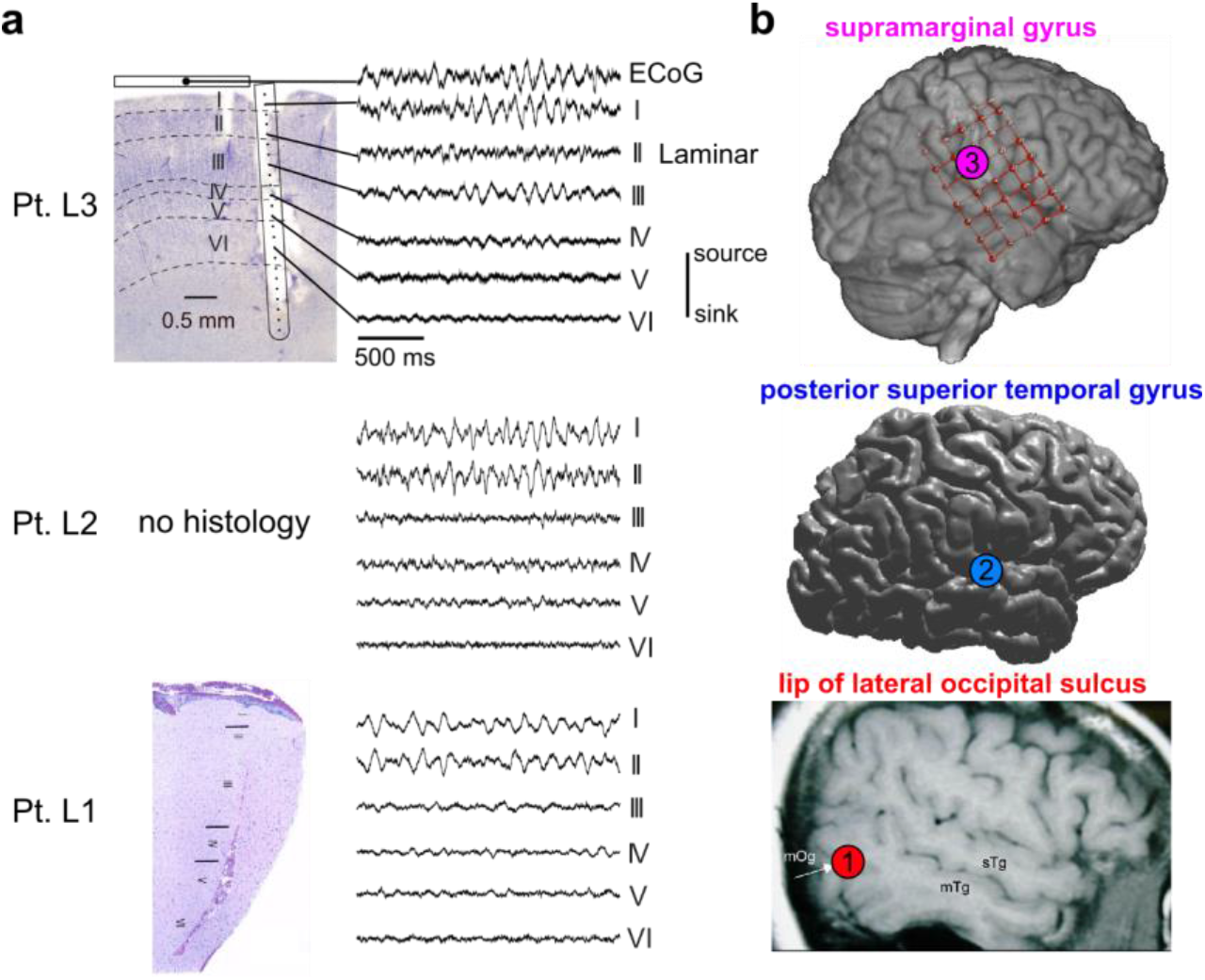
Laminar recordings of cortical alpha. **a** Nissl stains of the explanted tissue surrounding the laminar probe in patients L1 and L3, in addition to representative laminar traces from each layer in each subject. Note that despite being made in distinct cortical locations, alpha oscillations were always strongest in layer I/II. Furthermore, in L3 the trace of a simultaneously recorded overlying ECoG contact is near identical to the underlying laminar layer I. **b** Locations of each laminar probe in all patients.

**Fig. 6:**
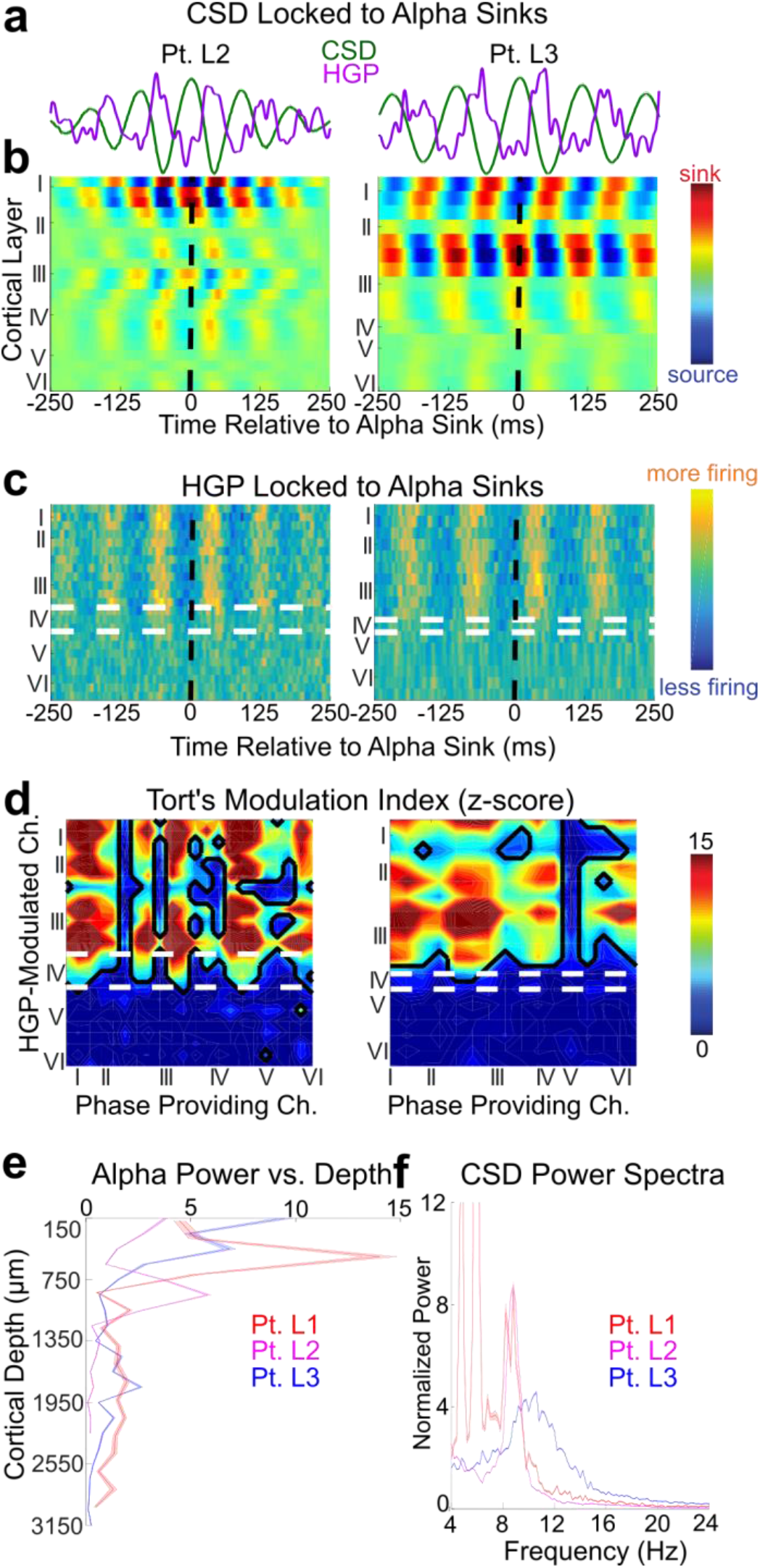
Alpha CSD and firing are maximal in supragranular cortex. **a** Average single-channel CSD and HGP waveforms of a single channel on same time-axes as **b** (±S.E.M. across alpha sinks). **b** CSD and **c** HGP averaged on current sinks in channels 3 and 6 in Pts. L2 and L3, respectively; white and black dashed lines indicate layer IV boundaries and the time of the alpha-sink, respectively. **d** Z-score of the MI between alpha phase and HGP across all channels. **e** Average alpha power throughout the cortical depth. (±S.E.M. across epochs). **f** Power spectra of the channel with greatest alpha power in each subject. (±S.E.M. across epochs).

Averaging HGP on alpha current sinks as well as measuring Tort’s MI between alpha CSD and HGP (**Fig. 6b-d**) reveals that alpha-band firing is located in layers I-III. Interestingly, while HGP was modulated by alpha throughout layers I-III significant MUA modulation was restricted to layer III (**Fig. 7**). It’s not clear whether this reflects differential sensitivity to noise, or suggests that MUA and HGP have divergent neural generators. However, both measures imply that the firing of supragranular (and not infragranular) pyramidal cells are phasic with the human alpha rhythm. Furthermore, this supragranular firing was maximal during very superficial sinks and minimal during superficial sources, consistent with active synaptic and/or voltage gated currents in layers I/II. The sink-over-source current dipole associated with increased firing would be recorded with ECoG/(M)EEG as surface-negative, as the negative end of the dipole (a sink of current flowing away from the extracellular space) is closer to the surface electrode. Consequently, this sink-source configuration comports with previous studies^37,40^ reporting that firing is maximal during the surface-negative trough of the alpha-rhythm and maximal during its surface positive peak.

**Figure 7:**
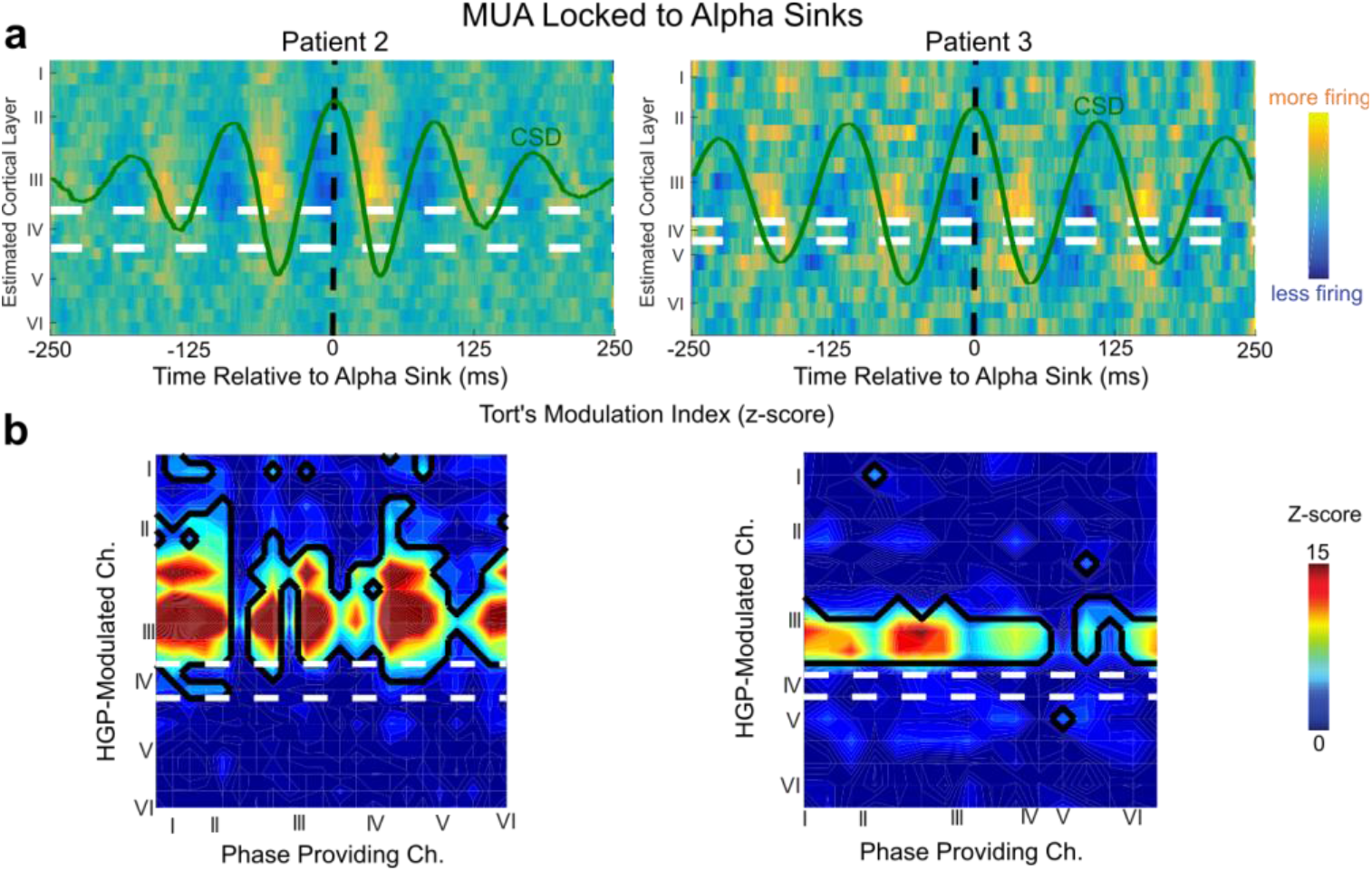
MUA is modulated in layer III. **a** MUA averaged on current sinks on the channels with the greatest alpha power (channels 3 and 6 respectively), which is most clearly modulated within lower layer III. **b** Tort’s Modulation Index between alpha CSD phase and MUA amplitude over all laminar contact pairs - note that firing is correlated with alpha phase in both superficial and deep cortex.

Lastly, we tested if the travelling alpha waves recorded with ECoG corresponded to the supragranular alpha oscillations recorded with laminar probes. To demonstrate this, we made combined ECoG-laminar recordings in Pt. L3 to record the same alpha rhythm at both local (laminar) and global (ECoG) scales^41^ (**Fig. 8**). Robust alpha oscillations (likely the classical mu rhythm given the electrodes’ locationsr^42^) were measured with ECoG from perirolandic cortex and with a simultaneously implanted laminar probe from the supramarginal gyrus; both recordings exhibited nearly identical alpha peaks (**Fig. 8e**), and averaging the laminar CSD on ECoG alpha troughs revealed strong supragranular sinks and sources (**Fig. 8f**). Alpha propagated as a travelling wave > throughout the grid (**Fig. 8a-d**) with a similar median speed (.62 m/s) to travelling waves in posterior cortex. If the supragranular alpha measured with laminar probes underlies the travelling alpha waves recorded with ECoG, we should be able to observe travelling alpha waves propagating through the supragranular layers of the laminar probe with a phase between its neighboring ECoG contacts (**Fig. 8c**). To establish this, we measured the average phase mean phase and found a phase lag intermediate to that of neighboring ECoG contacts (**Fig. 8a**, 335 this was also demonstrated by measuring the phase of the coherence between laminar and ECoG 336 in **Supplementary Fig. 10**). Interestingly, the direction of propagation was posterior-to-anterior 337 (**Fig. 8d**). While this may appear to conflict with our recordings of posterior alpha (which 338 propagated towards the occipital pole), the reversed propagation direction is consistent with 339 alpha propagating from higher to lower order cortex in both systems, as associative (higher-340 order) somatosensory areas are posterior to primary somatosensory cortex^43–45^.

**Fig. 8:**
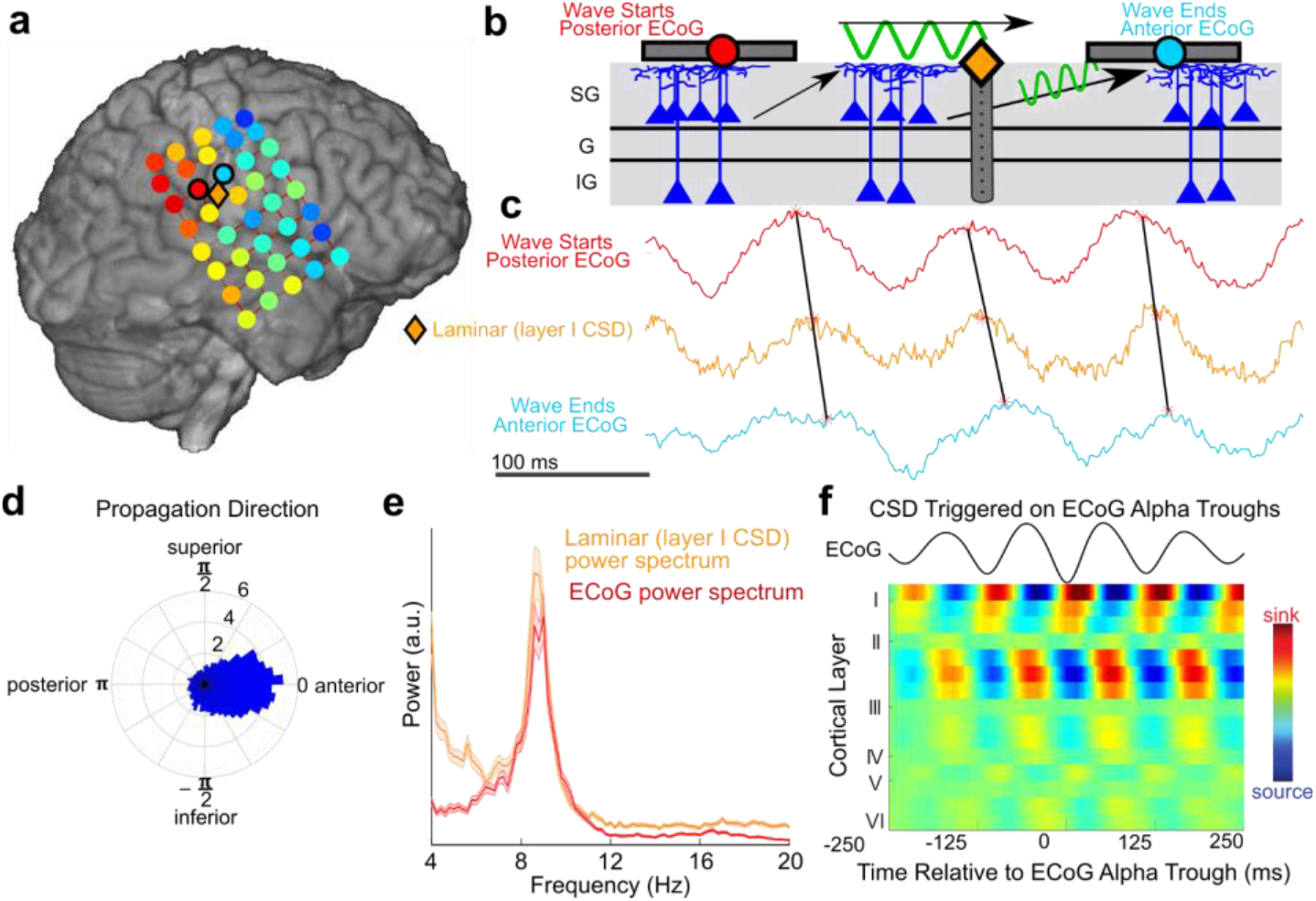
Simultaneous ECoG-Laminar recordings reveal travelling alpha waves which propagate through supragranular cortex. **a** Average circular distance of each ECoG (circles) and layer I laminar (diamond) contact’s alpha-phase from the spatial mean phase throughout the ECoG grid. Note that the laminar’s alpha phase is intermediate to neighboring ECoG contacts, suggesting that ECoG and the laminar probe recording the same travelling wave at different scales. **b** Representative drawing of a travelling alpha wave (as measured with ECoG) propagating through superficial layers (as measured by a laminar probe). **c** Example traces from ECoG contacts anterior (red) and posterior(blue) to the laminar probe. Alpha phase in laminar is intermediary to the ECoG contacts. **d** Distribution of travelling wave directions; mu waves propagate from posterior (higher-order) towards anterior (lower-order) cortex. **e** Laminar CSD averaged on troughs in the nearest ECoG contact. Note that alpha activity is superficial. **f** Power spectra from simultaneous laminar and ECoG recordings; note that they share a near identical alpha peak.

## Discussion

Our results suggest that alpha contributes to feedback processing within and across brain regions and structures (**Fig. 9**). The anatomical propagation of posterior alpha travelling waves from anterosuperior to posteroinferior cortex implies a functional progression from higher-order to lower-order visual areas, matching alpha’s putative role as a feedback rhythm^12^. Interestingly, previous scalp studies of human travelling alpha waves have found varying propagation directions^2^ in contrast to the consistent anterior-posterior directionality we observed. This may be due to propagation direction changing with task or behavioral state; our macaque recordings provide some evidence for this, as eye opening induced a clear (though smaller) peak of directional propagation from posterior to anterior areas (**Supplementary Fig. 4**). Alternatively, scalp recordings of alpha travelling waves may reflect a volume-conducted mixture of travelling alpha waves with different directions traversing distinct cortical hierarchies, consonant with our demonstration of posterior-to-anterior alpha propagation in somatosensory cortex. A study which sidestepped this concern by using ECoG didn’t claim to find the consistent propagation direction we did, though three of their four patients had broadly anterior-to-posterior directions. It should also be noted that they only reported linear propagation within a row/column of channels, rather than propagating throughout the entire grid as we find here. Alpha travelling waves might serve as a mechanism to internally scan the attentional field, tag distinct visual features with different phases^46^or facilitate plasticity between upstream and downstream areas^47,48^.

**Fig. 9:**
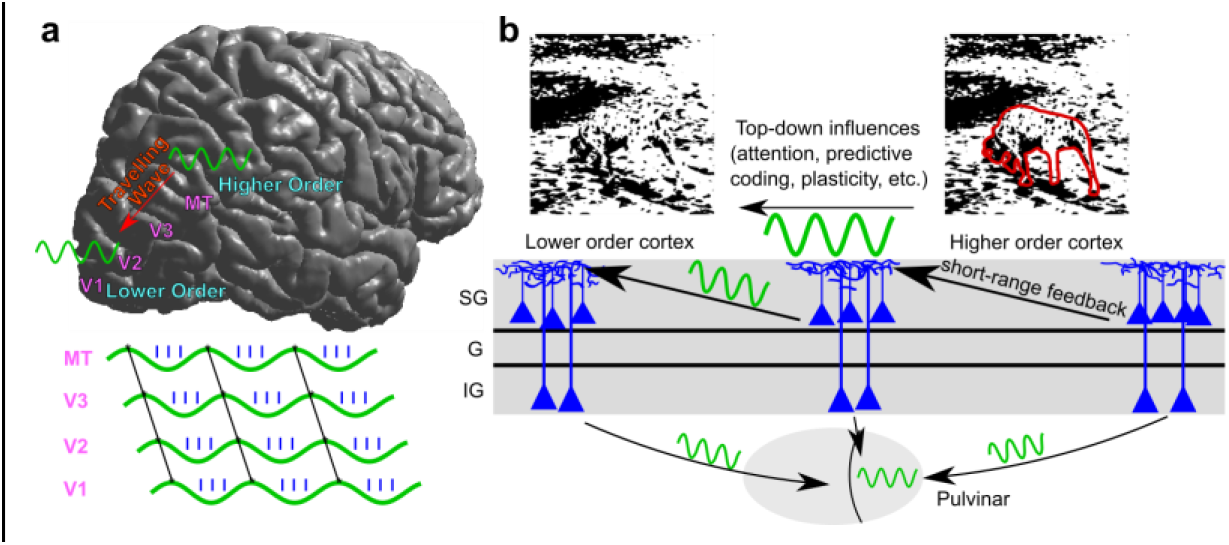
Alpha’s physiology could mediate feedback. **a** Alpha propagates as a travelling wave from higher order towards lower order cortical areas. **b** Alpha is strongest within supragranular cortex and may carry top-down information via short-range feedback connections to constrain lower-level processing. Cortical alpha in layer VI might influence alpha activity within the pulvinar.

Simultaneous recordings from human cortex and pulvinar provided several measures suggesting that cortical alpha leads thalamic alpha during quiet wakefulness. Thalamic alpha was less common than cortical alpha, and the cross-covariance of thalamic and cortical alpha power indicated that cortex (on average) led the thalamus. Furthermore, thalamic alpha LFPg was synchronous with cortical HGP (firing) more often than thalamic HGP; when thalamic alpha was synchronous with both cortical and thalamic HGP, HGP in the cortex led HGP in the thalamus. Finally, thalamic alpha was Granger-Caused by cortical alpha.

A potential weakness of these bivariate causal analyses is the possibility of a third structure (not recorded from) driving alpha oscillations in both pulvinar and cortex, such as the LGN^9^. While this is a possibility, the LGN is an unlikely cortical pacemaker as its major projections are limited to striate and circumstriate cortex^22^ and we found that alpha oscillations propagate towards (and not from) the occipital pole. Because the LGN and pulvinar are the two thalamic nuclei with the most robust projections to posterior cortex capable of driving visual alpha, these findings agree with strong cortical influences on thalamic (pulvinar) alpha during quiet wakefulness. Though this appears to contradict animal studies in which the pulvinar drove cortical alpha^4^, it’s consonant with findings that the cortex can still generate alpha in vitro^13^ and actually shows increased alpha-band power when the pulvinar is inactivated^23^, as well as alpha coherence within the cortex exceeding thalamocortical alpha coherence ^49^ These findings can be reconciled with other studies supporting thalamic alpha generators^4,50^ in a number of ways. There may be separable thalamic and cortical alpha pacemakers which become differentially active and coupled under different behavioral conditions^4^; the oscillatory circuit required to generate alpha may require both thalamic and cortical cells, or the pulvinar could enable an intracortical alpha pacemaker with tonic (nonrhythmic) excitation/inhibition without being a direct pacemaker^51^, perhaps coordinating cortical alpha phase in task-relevant fashion^4^. Though causal manipulations of alpha activity in animal models are needed to confirm this, the simplest explanation for our findings is a leading role for the cortex driving alpha in the pulvinar during quiet wakefulness. Functionally, corticothalamic alpha might inhibit the thalamus to gate feedforward processing and suppress irrelevant neural assemblies akin to its putative role within the cortex^7^, as low-frequency corticothalamic activity can inhibit thalamic firing^52^.

Laminar microelectrode recordings demonstrate that alpha oscillations reflect layer I/II currents (postsynaptic^36^) and layer I-III firing (presynaptic), demonstrating that supragranular layers are the source of alpha LFPs and HGP recorded via ECoG and (M)EEG. However, inferring the neural circuit mechanisms which generate the alpha rhythm from our laminar recordings is more complex. As supragranular pyramidal cells (which is where observed alpha-phasic firing) are known to make feedback projections to layers I/II (where we recorded driving currents), our recordings support layer II/III pyramidal cells as the primary alpha generators within the cortex during quiet wakefulness. Our ECoG recordings also support this, as the short range feedback projections subserved by supragranular pyramidal cells^53^ are a likely intracortical mechanism for mediating the continuously propagating top-down travelling waves measured using ECoG. These projections would enable oscillations which propagate continuously from high to low order cortex (i.e. not in the saltatory manner that might be expected if mediated by long-range feedback) at < 1 m/s (the conduction velocity of intracortical fibers^54^).(**Fig. 9**). While most models posit that layer V (infragranular) pyramidal cells drive alpha within the cortex^12,13^, these are based mostly on monopolar LFP recordings^37^ which (unlike CSD) are prone to volume conduction from deep sources. Some studies in macaque cortex appear to circumvent this by reporting significant alpha-band CSD-MUA coupling in deep layers of V1, V2 and V4^55^. However, it should be noted that these studies did not report CSD alpha power across the cortical depth (unlike another study which found that supragranular cortex had the most alpha power in each macaque primary sensory area^37^), and prior to calculating their directional measures they aligned their data to peaks/troughs in the channel with the most monopolar LFP alpha power. As this channel was likely infragranular (volume conduction leads to monopolar alpha power being spuriously maximal in deeper cortex^37^), this may have biased their results towards granular/infragranular generators. Importantly, CSD alpha power being greatest in superficial layers is consistent with alpha reflecting currents on the apical dendrites of supragranular or infragranular pyramidal cells; our paper resolves this ambiguity by measuring the coherence between alpha currents and multi-unit-activity throughout the cortical depth, and finding significant modulation of only supragranular firing by alpha currents (though a study in macaques did report modulation of granular MUA by superficial alpha CSD^37^). Further work employing causal manipulation of infragranular cortex in animal models will be needed to determine the role of deeper layers in alpha generation. In all, our microelectrode recordings strongly suggest that cortical alpha reflects short-range intracortical feedback mediated by supragranular pyramidal cells within superficial layers.

This supports an integrative function for alpha, due to the termination of widespread associative connections in superficial layers^53^ and the modulatory role of layer I/II apical dendrites^56,57^. A supragranular origin for the alpha rhythm is also in accord with its putative role in neural inhibition^58^, as layers I/II contain a dense interneuronal network which strongly inhibits the apical dendrites of excitatory cells throughout the cortical column^59^. This short-range inhibition would allow higher-order cortex to modulate the gain of lower order areas throughout visual cortex, providing a laminar circuit for top-down processes such as attention. Further studies which combine cognitive tasks with invasive recordings are needed to understand the implications of our findings for alpha’s behavioral role, as the physiology we describe is consistent with a breadth of potential functions for alpha. In all, we find that alpha acts within the nervous system by propagating from cortex to thalamus and higher-order to lower-order cortex, likely via short-range supragranular feedback projections. These intracortical and corticothalamic dynamics could allow alpha to sculpt activity throughout the neural hierarchy.

## Methods

### Patients

Implantations were performed on patients with pharmacologically-resistant epilepsy undergoing surgery to locate and resect seizure foci. Laminar and ECoG recordings were made from hospitals in the United States and Hungary, and thalamocortical depth recordings were performed in France. Seventeen patients (10 female, ages 15-50) were informed of potential risks and told that they had no obligation to participate in the study, as well as being informed that their decision to participate wouldn’t affect their clinical care. Experiments were made with fully informed consent as specified by the Declaration of Helsinki and approved by local institutional review boards. These boards included the Partners Health Care IRB, NYU Medical Center IRB, Stanford IRB, and the Hungarian Medical Scientific Council. All decisions concerning macroelectrode placement were made solely on a clinical basis, whereas laminar microelectrodes were inserted into cortex likely to be resected.

Patients were numbered according to their modality (E# for ECoG, S# for SEEG/macroelectrode depth and L# for laminars). Numbering for patients was started anew for each measurement modality, and no patients had more than one kind of electrode (ECoG, SEEG or laminar) analyzed with the exception of L3 (no corresponding ECoG number).

All recordings other than those during our eye-closure task were made of spontaneous activity during quiet wakefulness, in which the patient was not engaged in a cognitive task.

### General Analysis Procedures

Recordings were analyzed using custom MATLAB scripts with the CircStat^60^ and Fieldtrip^61^ Toolboxes.

Prior to further analysis, the raw data was visually inspected for artifacts due to machine noise, patient movement, or epileptiform activity. Epochs containing these artifacts (as judged by an expert neurologist) were removed prior to further analysis.

Unless otherwise specified, all analyses of alpha-band effects refer to the 7-13 Hz band. Error bars correspond to the standard error of the mean (S.E.M.).

Power and cross-spectral densities were found via the multi-taper method. This was performed by applying a Hanning taper and then taking the Fourier Transform of the zero-meaned data.

Coherence (**Fig. 4e, Supplementary Fig. 7a**) was calculated using the ft_connectivity function, which defines the coherence between mean subtracted time series x and y as 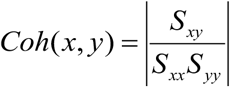, where *S_xy_* is the cross-spectral density between x and y and is the *S_xx_* autospectral density of x^62^. In order to quantify alpha-high gamma phase-amplitude-coupling, we measured the coherence between the broadband LFP and Hilbert amplitude of the high-gamma filtered data. To determine the statistical significance of coherence between all channel pairs, we generated a reference distribution of coherence values under the null hypothesis of no temporal relationship between each pair of time-series^63^. This was accomplished by shuffling the temporal order of 2-second epochs for each channel 200 times, and then recalculating the coherencies between all channels. To calculate the significance of coherencies in the alpha-band, we summed the coherence between 7-13 Hz for each permutation, and then used the mean and standard deviation of this reference distribution of alpha coherencies to determine the z-score of the real summed coherence between 7-13 Hz. Coherencies were deemed significant at p < .05, Bonferroni corrected within patients (the critical value being p < .05 / (number of channel pairs) for each subject).

To derive alpha and high-gamma amplitude as well as alpha phase, we used the Hilbert Transform. First, data was filtered using a two-pass fourth-order IIR Butterworth Filter. Then, the analytic signal *z*(*t*) was found by applying the Hilbert Transform to the filtered signal of each channel. The phase series *φ*(*t*) was found by taking the angle of the analytic signal, and the amplitude *A*(*t*) of every channel was found by taking the real component of the analytic signal.

To determine the effects of alpha rhythms on neural firing, we used Tort’s Modulation Index^27^ (**Fig. 4f, Supplementary Fig. 7b**) with a non-parametric trial shuffling procedure to assess significance. First, we applied the Hilbert Transform (see above) to derive amplitude series *A*(*t*) and phase series *φ*(*t*). *φ*(*t*) was then reordered from – *π* to + *π*, and *A*(*t*) for every other channel and frequency was reordered using the same permutation vector. Amplitude was then averaged within 36 bins of phase (i.e. 10 degrees) and normalized by the sum over bins, yielding *φ*. The modulation index (MI) was then calculated as 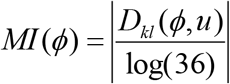 for each channel and frequency pair, where *D_kl_* is the Kullback-Leibler divergence, *u* is the uniform distribution (i.e. no relationship between amplitude and phase) and log(36) is the natural logarithm of the number of phase bins^27^. *D_kl_* was computed as log(36) – H(P), where H(P) was the distribution’s Shannon’s Entropy.

Statistical significance of MI values was determined similarly to coherence (a reference distribution under the null hypothesis of a random relationship between amplitude and phase was formed, the MI was calculated for each shuffled data set and the mean and variance of these null hypothesis derived MIs at each channel and frequency were used to determine the z-score of the actual MIs at each channel and modulating/modulated frequency pair). Instead of shuffling the temporal order of two second epochs to create surrogate datasets we iteratively split the phase series into two epochs 200 times, the split point being 20-80% through the length of the data, and swapped their order.

### Travelling Waves

We utilized 4.54±.87 minutes (mean ± standard deviation) of ECoG recordings made for clinical purposes. These arrays had 2-mm contact diameters and 1-cm intercontact spacing, and were referenced to 1-4 inactive electrodes placed outside of the dura facing the skull. In Patients 1, 3 and 4, patients were instructed to open and close their eyes with an audial cue at 15 second intervals using Presentation software (Neurobehavioral Systems, Albany, CA, USA). We only utilized activity during eye-closure in these patients (except for **Supplementary Fig. 4**). In Patients E2 and 5 (who didn’t participate in the eye-closure task), we analyzed spontaneous activity during quiet wakefulness. We also analyzed 16.5 minutes of open-source ECoG recordings made from a macaque monkey during an eye closure task. Eyes were closed via a sleep mask for 10 minutes, and the sleep mask was then removed for 10 minutes of data. The Macaque was included in group statistics with other patients due to its high similarity with human activity (**Supplementary Fig. 4**). Further details concerning the macaque recording can be found at Neurotycho.org^20^. Time-domain data in Pts. E1 and E5 were spatially interpolated in missing channels (using inpaint_nans^64^) prior to further analysis.

To localize contacts to the pial surface, we aligned a pre-operative MRI with a structural MRI or CT. These contact locations were then displayed on the reconstructed cortical surface, created using Freesurfer^65^, of each individual patient (**Figs. 1-2**) ^66,67^.

This analysis was performed separately for strips and grids as well as strips >2 cm apart, as the large cortical distances between them would make phase differences difficult to interpret.

We then wished to measure the directionality of these travelling waves. To do this we employed the phase gradient ∇, found by using MATLAB’s gradient function (but with subtractions replaced with circular distances). To prove that there was a consistent directionality of propagation across time, at each time-point we found the mean direction of the gradient throughout the grid. Using Rayleigh’s test for non-uniformity demonstrated that each patient had a significant propagation direction (**Supplementary Fig. 4**). To generate **Fig. 2d**, we binned the travelling wave directions across time into 100 bins normalized within patients (i.e. divided the count of each bin by the total number of time-points), and then averaged across patients (with the error bar being the S.E.M. across patients).

To find instantaneous speeds across time, the grid’s instantaneous frequency first needed to be estimated. This was done by filtering from 7-13 Hz with a plateau-shaped filter with a transition width of 15%. A plateau-shaped filter was used instead of a more traditional Butterworth or FIR because the former isn’t Gaussian in the frequency domain, and therefore does not bias the instantaneous frequency towards the center of the band-pass range^68^. The phase of the band-pass filtered data was then estimated using the angle of the Hilbert transform, and subsequently unwrapped. The first derivative of this phase series was defined as the difference in phase between consecutive samples, using MATLAB’s diff function. Finally, to attenuate the sensitivity of this technique to noise, a median filter with a kernel of 10 samples was used^68^.

Instantaneous speeds were calculated as follows: each channel’s instantaneous frequency (see above paragraph) was divided by the magnitude of its phase gradient 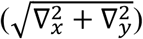, yielding the instantaneous speed at each channel and timepoint. Then, at each timepoint, the median speed and frequency across all channels was found, and timepoints with a speed or frequency in the top or bottom .5^th^ percentile was rejected to eliminate outliers. The distribution of these median speeds across time was then plotted as a normalized histogram (bin width of .01 m/s) for each patient and presented **Supplementary Fig. 4**. These normalized histograms were averaged and plotted in **Fig. 2c**.

To ensure that this effect was specific to the alpha-band, we re-applied our main analysis to 2-Hz filtered bands with 1 Hz spacing from 1 to 35 Hz and found the resultant vector length of propagation direction across all time points. A clear alpha-peak is observed (**Supplementary Fig. 3**).

### Corticothalamic Interactions

Stereoencephalography (SEEG)^69^ was performed on 9 patients to characterize epileptogenic activity and inform possible resections. SEEG macroelectrode depth probes had 10-15 contacts; each contact was 2 mm long and .8mm in diameter with 1.5mm inter-contact spacing. The probes themselves were ~5 cm long, with the exact length varying between electrodes. In Pts. S1-7, contact locations were found by stereotactic teleradiographs from within the stereotactic frame. These coordinates were then superimposed on a T1 MRI of the subject. An atlas^70^ was then overlaid to determine the anatomical positions of thalamic and cortical contacts (**Fig. 3a**). Contacts were localized in Pts. S8-9 by aligning a post-operative CT with a pre-operative MRI. Adjacent channels (when at least one was in the grey matter) were referenced to each other to assure local generation of measured activity. Further use of contact, channel or site all refer to these bipolar channel pairs. Recordings were made at 256 Hz in Pts. S1-3, 128 Hz in Pts. S4-7 and 1024 Hz in Pts. S8-9. For further details see ^71^.

We analyzed spontaneous activity (36 ± 7.5 min., mean±standard deviation) during wakefulness prior to the onset of sleep, the time of which was determined behaviorally as well as electrographically by a qualified sleep-stager using standard methods^72^. The last three minutes of wakefulness before the onset of sleep was rejected to further avoid the analysis of sleep or excess drowsiness.

Prior to further analyses, we split the data into non-overlapping two-second epochs. We bandpass filtered (two-pass 3^rd^ order Butterworth) thalamic activity in the alpha-band, then found the absolute value of its Hilbert transform to find single-trial thalamic alpha amplitude. Then, the 20% of epochs with the most total alpha-band amplitude (summing across thalamic bipolar pairs and samples) were used for further processing.

Cortical and thalamic power spectral peaks were found using the peakfinder^73^ algorithm with a selectivity (the minimum difference a local maxima must have from the nearest local minima to be considered a peak) of the power spectrum’s range divided by 5. A channel was considered to have an alpha peak if it had at least one peak between 7-13 Hz.

To phase-align ongoing alpha activity (**Fig. 4d**), we picked the thalamic contact with the greatest alpha-band power and averaged the rest of our data (cortical and thalamic wide-band LFP and HGP) to alpha-band peaks in this channel. Alpha band peaks were found by bandpass filtering from 7-13 Hz (two-pass 3^rd^ order Butterworth), taking the angle of the Hilbert Transform to find the phase, and then finding peaks in this series. LFP was high-pass filtered at 2 Hz (two-pass third-order Butterworth) prior to averaging on alpha peaks.

High-gamma-power (HGP) was derived by filtering from 70-120 Hz in Pts. S1-3 (due to a Nyquist frequency of 128 Hz) and 70-190 Hz in Pts. S8-9. The sampling rate in Pts. S4-7 (128 Hz) was too low to measure HGP. Although the former frequency band is somewhat lower than the usual definition of HGP (70-190 Hz), we observed similar results in patients with both usual and reduced HGP bands (**Supplementary Fig. 7**). Furthermore, a previous study employing the same recordings demonstrated that 70-120 Hz HGP reliably decreased during K-complexes identical to 70-190 Hz power^71^. Lastly, reanalyzing Pts. S8-9 using the 70-120 Hz band yielded highly similar results. 7/14 (coherence) or 10/14 (MI) thalamic channels exhibited significant PAC between thalamic alpha LFPg and the HGP of at least one cortical channel (compare to 9/14 coherence and 10/14 MI with 70-190 Hz band). Furthermore, 0/14 (coherence) or 4/14 (MI) thalamic channels had significant PAC between thalamic alpha LFPg and thalamic HGP (0/14 (coherence) and 3/14 (MI) with the 70-190 Hz band).

The Sharpness Ratio (SR) was measured as described in Cole 2017^29^. Briefly, the steepness of each extremum in the alpha-band filtered data was calculated as 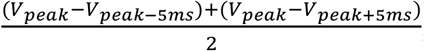, where *V_peak_* and *V*_*peak*±5*ms*_ denotes the raw voltage at each alpha-band extremum and 5ms before or after, respectively. The average ratio of trough to peak sharpness and vice-versa was then calculated, with the SR for a given channel being the greater of the two ratios.

Granger Causality (GC) analyses were performed using the Multivariate Granger Causality Toolbox ^32^. Frequency-resolved GC values were assessed between each corticothalamic channel pair in the non-overlapping 2-second epochs with the greatest 20% of thalamic alpha amplitude as found by the Hilbert Transform. In order to measure causality in the alpha-band without using a prohibitively large model-order, we then downsampled our data to 256 Hz and used a model order of 26 across all channels and patients (13 in patients with 128 Hz sampling rate). To ensure stationarity, we high-pass filtered at .2 Hz (Butterworth 2nd order two-pass) and then demeaned, detrended and z-scored each channel within each epoch. The lag needed for the autocovariance sequence of each epoch’s vector auto-regressive model (VAR) to decay below the default numerical tolerance (10^−8^) was computed with autocov_to_pwcgc(). Epochs which had an autocovariance lag of at least 2000 samples were deemed inappropriate for VAR modelling and were not used for Granger analysis. Conceptually, it would be most interpretable to measure GC between the LFPg and HGP in thalamus and cortex to determine whether thalamic currents/firing modulate cortical firing/currents (or vice-versa). Unfortunately, this is precluded by the deleterious effects of filtering on GC, as the filtering necessary to measure HGP can seriously degrade GC, create spurious causality and even reverse its directionality^74,75^.

### Laminar Recordings

Laminar microelectrode arrays were inserted on the basis of two criteria: first, the tissue must be very likely to be resected^76^. This could be because it was clearly within the seizure onset zone, or because it was healthy tissue overlying the seizure onset zone which would have to be removed during the resection. Secondly, the cortex in question must have had no chance of being eloquent. In all three patients, the tissue surrounding the laminar probe was ultimately resected. A silicone sheet attached to the array’s top was used to keep the probe perpendicular to the cortical depth, with surface tension between the sheet and the pia, as well as pressure from the overlying grid and dura, keeping the array in place. This sheet also ensured that the laminar array was perpendicular to the cortex and that the first contact was placed just below the cortical surface (depth of ~175 microns in Pt. L1 and ~150 microns in Pts. L2-3). Each laminar probe spanned the cortical depth with a length of 3.5 mm and diameter of .35 mm. Contacts had 40 micron diameters, spaced at 175 microns in Pt. L1 and 150 microns in Pt. L2-3. Recordings were made during 11.32 ± 0.48 min. (mean ± standard deviation) of quiet wakefulness.

The local-field-potential-gradient (LFPg), or the first derivative of the field potential (i.e. each contact referenced to its neighbor) and multi-unit activity (MUA) were recorded simultaneously at 2000 and 20000 Hz and filtered online from .2-500 Hz and 200-5000 Hz, respectively. Data from faulty channels (2 in each patient) were linearly interpolated from the channels directly above and below them.

Line noise was eliminated from both the LFPg and MUA by band-stop filtering (4 Hz bandwidth) at 60 Hz in Pts. L1-2 and 50 Hz in Pt. L3 (4th order Butterworth). The LFPg was then high-pass filtered at .5 Hz in Pts. L2-3 and 3.5 Hz in Pt. L1 due to a low-frequency vascular artifact (two-pass 2nd order Butterworth), HGP from 70-190 Hz and MUA from 300-3000 Hz (4th order Butterworth). MUA was then rectified and resampled at 2000 Hz. The data was then sub-sampled into two-second artifact-free epochs and, consistent with ECoG and SEEG, the 20% of epochs with the most alpha-band CSD amplitude across all channels was utilized for further analysis. In Pt. L1, all artifact-free epochs (not just those with the most alpha amplitude) were used due to a relatively short recording session. We observed no significant modulation of HGP or MUA in Pt. L1, probably due to technical issues with the recording such as gliosis or faulty electrodes.

Current source density (CSD) was measured by taking the first spatial derivative of the LFPg (in effect the second spatial derivative of the monopolar field potential) and then applying a 5-point Hamming filter^33,34^. The Vaknin approximation (adding pseudo-channels of zeros to the LFPg above and below the array) was used to estimate the CSD on the second most deep and superficial channels of the laminar probe^77^.

To visualize the profile of alpha CSD and HGP, we picked the laminar contact with the highest average alpha power, and averaged our HGP and CSD recordings to alpha-band peaks in the CSD of this channel. Alpha band peaks were found by bandpass filtering from 7-13 Hz (two-pass third-order Butterworth), taking the angle of the Hilbert Transform to find the phase, and then finding peaks in this series (corresponding to alpha-band current sinks). Average MUA displays were then smoothened with a Gaussian filter for display purposes (width of 5 ms and 750 microns, *σ_x_*=10 ms and *σ_y_*= 112.5 microns).

Prior to visualizing laminar power spectra as well as alpha power across channels (**Fig.6e-f**), power spectra for each subject were normalized by dividing by the mean power from 4-25 Hz across all channels and epochs.

We localized laminar contacts to cortical layers by performing histology on explanted tissue in Pts. L1 and L3, and identifying a putative layer IV sink in Pt. L2 (**Supplementary Methods**)

### Data Availability

We will make data available to the degree it is possible upon request given participant consent restraints and HIPAA requirements. Macaque data is publically available at http://neurotycho.org/data/20120813ktanesthesiaandsleepchibitoruyanagawa.

### Code Availability

Code will be made available from the corresponding author upon reasonable request.

## Acknowledgements

The authors thank Erica Johnson, Nathan Meng, Richárd Fiáth, and Adam Niese for insightful comments, hypotheses and technical support. We also thank Project Tycho for making their macaque dataset publically available. This study was supported by the U.S. Office of Naval Research Grant N00014-13-1-0672, National Institutes of Health Grants R01-MH-099645, R01-EB-009282, R01-NS-062092, K24-NS-088568 and the MGH Executive Council on Research, Hungarian National Brain Research Program grant KTIA_13_NAP-A-IV/1-4,6, EU FP7 600925 NeuroSeeker, and Hungarian Government grants KTIA-NAP 13-1-2013-0001, OTKA PD101754, OTKA K119443.

## Author Contributions

S.S.C., I.U. and E.H. conceived of the experiments; R.M.M, L.W., L.E., O.D., W.D., H.B., MR., P.C., I.U., D.F., G.H., E.E, and S.S.C. conducted the experiments; M.H. analyzed the data; M.H., E.H., A.M. and S.S.C. interpreted the results; M.H., and S.S.C. wrote the manuscript, and all authors discussed and edited the manuscript.

